# Dining in the daylight: Nocturnal rhinoceros beetles extend feeding periods on host trees with reduced sap exudation

**DOI:** 10.1101/2023.12.25.573284

**Authors:** Asahi Kanda, Ryo Shibata, Takahiro Ueno, Namifu Yamashita, Wataru Kojima

**Author notes:** **Correspondence:** Asahi Kanda, Graduate School of Sciences and Technology for Innovation, Yamaguchi University, 1677-1 Yoshida, Yamaguchi City, Yamaguchi 7538511, Japan.

## Abstract

While the Japanese rhinoceros beetle *Trypoxylus dichotomus* typically feeds on the sap of the oak *Quercus acutissima* and the crape myrtle *Lagerstroemia subcostata* during the night, it exhibits feeding activity both during the day and night when it utilizes the ash tree *Fraxinus griffithii*. However, the mechanisms underlying the variations in temporal activity patterns remain unknown. We compared feeding rates (measured as body mass increments) and sap exudation rates among *F. griffithii, Q. acutissima*, and *L. subcostata*. We found that beetles feeding on *L. subcostata* and *Q. acutissima* exhibited significantly higher feeding rates than those feeding on *F. griffithii*. No significant differences in feeding rates were observed between *L. subcostata* and *Q. acutissima*. The sap exudation rate was significantly higher for *Q. acutissima* than for *F. griffithii*. However, there were no significant differences in the sap exudation rates between *F. griffithii* and *L. subcostata* or between *Q. acutissima* and *L. subcostata*. These findings suggest that lower feeding rates on *F. griffithii* prolong the feeding duration, resulting in daytime activity. While the low sap exudation in *L. subcostata* seems inconsistent with high feeding rates on this host, this apparent contradiction could be related to the extended duration of sap exudation.

## Introduction

The daily activity patterns of animals have evolved through natural selection to maximize their fitness. Although activity patterns are typically fixed within species and are classified as diurnal, nocturnal, or crepuscular, some species exhibit adaptive changes in their activity patterns in response to various factors (Levy et al. 2018), including developmental stage (Paulissen 1988), predation pressure (Fenn and Macdonald 1995; Shiojiri et al. 2006), competitor (Siva-Jothy 1987; Levy et al. 2007), season (Stiles et al. 2017), temperature (Nisimura et al. 2005) and physiological state. Hunger is a primary physiological factor influencing animal activity patterns (Lockard 1978; Kramer et al. 2001; Pereira 2010; Hut et al. 2011). Starvation compels individuals to engage in foraging activities to secure the necessary energy, causing them to deviate from their original temporal niches, even if such behavior incurs some costs.

The Japanese rhinoceros beetle *Trypoxylus dichotomus* (Coleoptera, Scarabaeidae, Dynastinae) feeds on tree sap exudates. In Japan, beetles utilize various host tree species across different taxonomic groups, such as Fagaceae, Ulmaceae, and Oleaceae (Siva-Jothy 1987; Hongo 2006; Moriue 2009; Yagihashi et al. 2014; Del sol et al. 2021). Although the rhinoceros beetle is typically nocturnal, its activity patterns vary plastically depending on the host species. Temporal activity patterns were quantitatively investigated in two host species: the oak *Quercus acutissima* and the ash tree *Fraxinus griffithii*. In *Q. acutissima, T. dichotomus* feeds on sap from wounds formed by wood-boring insects, such as the carpenter worm *Cossus jezoensis* (Yoshimoto et al. 2005; Ichikawa and Ueda 2010), the bark beetle *Platypus quercivorus* (Akaishi et al. 2006), and the white stripe long-horned beetle *Batocera lineolate* (Takakuwa 2007). *T. dichotomus* flies to sap sites on *Q. acutissima* after sunset and leaves its roosts before sunrise; therefore, beetles are rarely seen during the daytime (Siva-Jothy 1987). In contrast, *T. dichotomus* carves the bark of *F. griffithii*, using its mandible to feed on the sap with head-scooping movements (Hongo 2006; Ichiishi et al. 2019). *T. dichotomus* individuals fly to *F. griffithii* trees at night and remain at their feeding sites for extended periods compared to their feeding behavior on *Q. acutissima*. Consequently, more than half of them continue to feed during the day (Shibata and Kojima 2021). Despite the distinct contrast in activity patterns among host species, the underlying mechanisms remain unclear.

Since wounds on *F. griffithii* formed by *T. dichotomus* exuded less sap than those on *Q. acutissima* (R. Shibata, personal observation), we predicted that lower feeding rates on *F. griffithii* would prolong feeding duration, resulting in daytime activity. To confirm this prediction, we compared the temporal activity patterns, sap exudation rates, and feeding rates of beetles among the host species. In addition to *F. griffithii* and *Q. acutissima*, we dealt with the crape myrtle *Lagerstroemia subcostata*, which is taxonomically distinct from the other two hosts.

## Materials and Methods

### Activity Patterns on Lagerstroemia subcostata

Although the activity patterns of *T. dichotomus* have been previously reported for two host species, *Q. acutissima* and *F. griffithii* (see Introduction), no such information is available for *L. subcostata*. Therefore, we assessed the activity patterns of beetles feeding on *L. subcostata*. The study was conducted from August 6 to August 10, 2021, along a path (total length of approximately 300 m) in Anbo, Yakushima-cho, Kagoshima Prefecture, Japan. Twenty-five trees were selected for observation based on preliminary surveys, and the number of beetles on these trees was counted hourly. Surveys were not conducted during rainy weather, and there were 83 counting events over the course of the study. If the beetles were high up in the trees, they were observed using binoculars (Atrek II HR 8 × 32 WP; Vixen). For night-time observations, a headlight (MM285-h; GENTOS) was used; however, direct illumination of the beetles was avoided.

### Feeding Rate

The weight gain of beetles per hour was measured as an indicator of feeding rate. The experiments were conducted in Sugito Town, Saitama Prefecture, in August 2021 for *F. griffithii* (n = 20 beetles); Yamaguchi City, Yamaguchi Prefecture, in August 2021 for *Q. acutissima* (n = 26 beetles); and Yakushima-cho in August 2022 for *L. subcostata* (n = 36 beetles). All measurements were conducted at night (20:00 ∼ 26:00) when *T. dichotomus* is most active. Prior to the experiments, field-collected beetles were fasted for 3 days to stimulate their feeding behavior. Beetles were weighed using a portable digital scale (SF-700; TUO) at an accuracy of 0.01 g. They were then immediately released into the trees. They typically initiated sap feeding within approximately 10 min. After an hour of feeding, beetles were collected and weighed. Individuals who did not continuously feed were excluded from the analysis. Beetles defecated during feeding, potentially influencing the accuracy of the feeding rate estimate. However, our experimental setup did not allow us to measure the weight loss caused by defecation.

### Sap Exudation Rate

The sap exudation rate at the beetle feeding sites was measured following the method described by Yoshimoto (2007). The experiments were conducted in Mitaka City, Tokyo, in July 2023 for *F. griffithii*; Atsugi City, Kanagawa Prefecture, in July 2023 for *Q. acutissima*; and Yakushima-cho in July 2023 for *L. subcostata*. As our aim was to test the relationship between activity time of beetles and sap exudation rates, we measured these rates during the beetles’ activity periods on the focal host species. Specifically, measurements were conducted at night (20:00 ∼ 24:00) for *Q. acutissima* (n = 5 trees and 10 wounds) and *L. subcostata* (n = 2 trees and 8 wounds), and both during the daytime (10:00 ∼ 14:00, n = 5 trees and 10 wounds) and at night (20:00 ∼ 24:00, n = 5 trees and 10 wounds) for *F. griffithii*. The beetles were removed from the feeding sites, and solids and liquids were wiped away with a paper towel. An unused paper towel (approximately 1.5 g), the weight of which was measured beforehand, was pressed against the wounds to absorb the sap. The paper towel was covered with cling wrap to prevent evaporation. After 20 min, the towel was collected and weighed again. The increase in weight was used as an indicator of the sap exudation rate.

### Statistical Analyses

Tukey’s multiple comparison method (HSD) was used to determine variation in feeding rates among the three host species. As our preliminary analyses showed that pronotum width (an indicator of body size) and sex did not significantly affect the feeding rate, these variables were not included in the model. Comparisons of sap exudation rates among host species were conducted using TukeyHSD. The R software package (version 4.0.4; R Foundation for Statistical Computing, Vienna, Austria) was used for statistical analysis.

## Results

The number of *T. dichotomus* on *L. subcostata* started increasing around 8 pm, reaching a maximum at midnight, and most individuals left the tree by 5 am. The number of beetles during the daytime was approximately one-fifth of the nighttime peak. Most of the individuals on the trees were engaged in activities such as feeding and mating. These results indicated that the beetles feeding on *L. subcostata* were nocturnal.

The weight gain of the beetles feeding on *L. subcostata* and *Q. acutissima* was significantly greater than those feeding on *F. griffithii*. There was no significant difference in the weight gain between those feeding on *Q. acutissima* and *L. subcostata*. The body weights of some individuals decreased after 1-hour feeding, probably because of defecation.

Since the sap exudation rate of *F. griffithii* did not vary between day and night (Welch’s t-test: t = -1.2358, df = 17.89, p = 0.2325), the data were combined for comparison with other tree species. The sap exudation rate of *Q. acutissima* was significantly higher than that of *F. griffithii*. There were no significant differences in the sap exudation rates between *F. griffithii* and *L. subcostata* or *Q. acutissima* and *L. subcostata*.

## Discussion

This study revealed that the feeding rate of *T. dichotomus* on *F. griffithii* was lower than that on other host species. In addition, the sap exudation rate of *F. griffithii* was lower than that of *Q. acutissima*. Furthermore, we found that the beetle activity patterns on *L. subcostata* were similar to those on *Q. acutissima*. Based on these findings, when beetles utilize host trees with low sap exudation, their activity periods are likely to be extended.

Feeding behavior on *Q. acutissima* and *L. subcostata* is probably completed during the night, whereas beetles feeding on *F. griffithii* face the challenge of inadequate sap intake at night, compelling them to continue feeding until daytime.

Sap quantity in *Q. acutissima* was greater than that of the other tree species. This may be due to the sap exudation mechanism of *Q. acutissima* was different from that of the other tree species. In *Q. acutissima*, the interior of the trunk was damaged by wood-boring insects, whereas in the other two tree species, only the surface of the bark was carved by *T. dichotomus*. There was no significant difference in sap exudation rates between *F. griffithii* and *L. subcostata*. Nevertheless, the feeding rates were significantly higher in *L. subcostata*. These results are seemingly contradictory but could be explained by the difference in the duration of sap exudation. *L. subcostata* may continue to produce sap longer with a single excavation than *F. griffithii*.

Diurnal activities are likely to impose costs on *T. dichotomus*. The large black body of *T. dichotomus* makes it more conspicuous during the day, increasing its vulnerability to visual predators such as crows (Kojima et al. 2014). Furthermore, the daytime ambient temperature in central Japan reaches 35–40 °C, which may cause physiological stress to *T. dichotomus*, as has been reported in other insect species (Chen et al. 2018). Despite these potential costs, *T. dichotomus* might gain net benefits from the extended activity period of *F. griffithii*.

Although this study suggests that feeding rate affects the activity patterns of rhinoceros beetles, multiple unexplored factors could also be attributable to host-dependent activity patterns. For example, insufficient nutritional content (e.g., sugar and protein) in the sap may lead to prolonged feeding times. Phloem sap consists mainly of sugars, with trace amounts of amino acids, organic acids, and proteins (Jensen et al. 2016). For example, the sap from *Q. acutissima* contains fructose, glucose, ethanol, acetic acid, glycerol, lactic acid, and 2,3-butanediol (Yoshimoto 2008). However, due to the absence of sap composition data for the other two host species, we cannot examine or test this hypothesis.

Interspecific competition for sap sites is also one of the potential factors that might affect the activity patterns of rhinoceros beetles. Kojima (2023) reported that the giant hornet *Vespa mandarinia* visited sap sites on *Q. acutissima* at approximately 5:00 am and physically excluded *T. dichotomus*. After the hornets were experimentally removed, the beetles continued to occupy the sap sites until noon. Thus, the activity duration of rhinoceros beetles may be extended into the daytime even on *Q. acutissima*, if they cannot obtain adequate sap due to factors such as lower conditions of sap sites or intense intraspecific competition. Additionally, male beetles are active at different times of the day depending on their body size (Siva-Jothy 1987). As the body size of individuals at the sap sites was not measured in this or previous studies (Shibata and Kojima 2021; Kojima 2023), further investigation of individual-level activity patterns is required.

Host-dependent variation in activity patterns has been observed in the sap-feeding stag beetle *Prosopocoilis dissimilis okinawanus* (Hongo 2005). While some individuals of this stag beetle exhibit daytime activity on *Citrus trichotomum* var. esculentum and *Persea thunbergii*, they become exclusively nocturnal with an adequate food in laboratory. Although the study did not examine the sap feeding rates and the sap exudation rates, similar mechanisms to those observed in *T. dichotomus* might underlie the activity patterns of the stag beetle. This example in the stag beetle and our study in *T. dichotomus* also highlight the possibility that host-dependent variation in activity patterns may be widespread across sap-feeding insects.

## Acknowledgements

We thank Kazuya Shibata and Mariko Shibata for assistance with field observation, and Yoshiaki Kobayashi for the permission to perform fieldwork and provision of information.

## Funding

This study was partly supported by grants from Yamaguchi University.

## Declarations Conflict of interest

The authors declare that they have no conflict of interest.

## Ethical approval

No approval of research ethics committees was required for this research.

---

**Figure 1.**
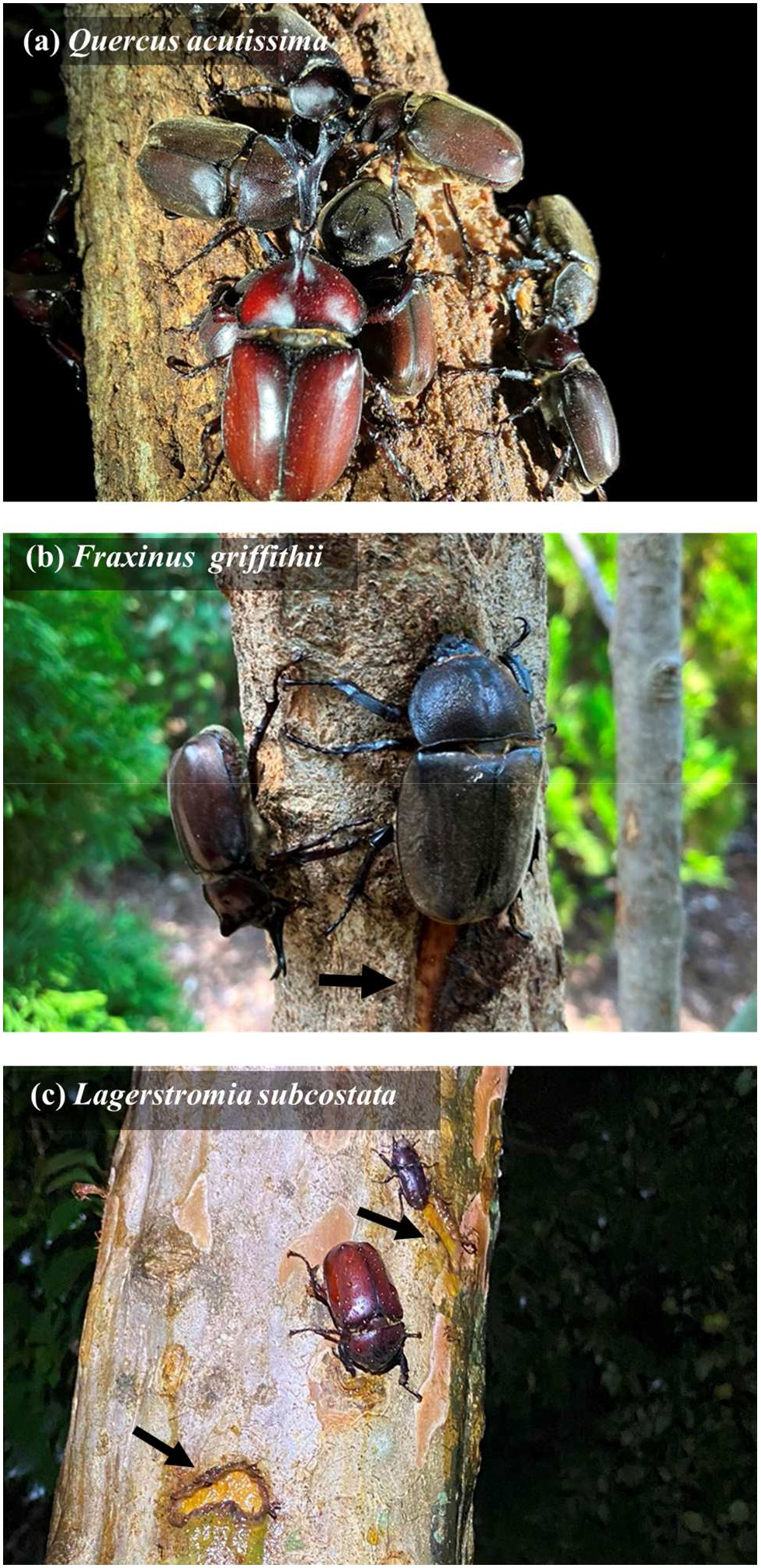
*Trypoxylus dichotomus* feeding on the sap of *Quercus acutissima* (a), *Fraxinus griffithii* (b), and *Lagerstroemia subcostata* (c). In (b) and (c), *T. dichotomus* is engaging in bark carving, with the scars indicated by black arrows. In (c), the smaller beetle above the female *T. dichotomus* is a female stag beetle *Prosopocoilus inclinatus*.

**Figure 2.**
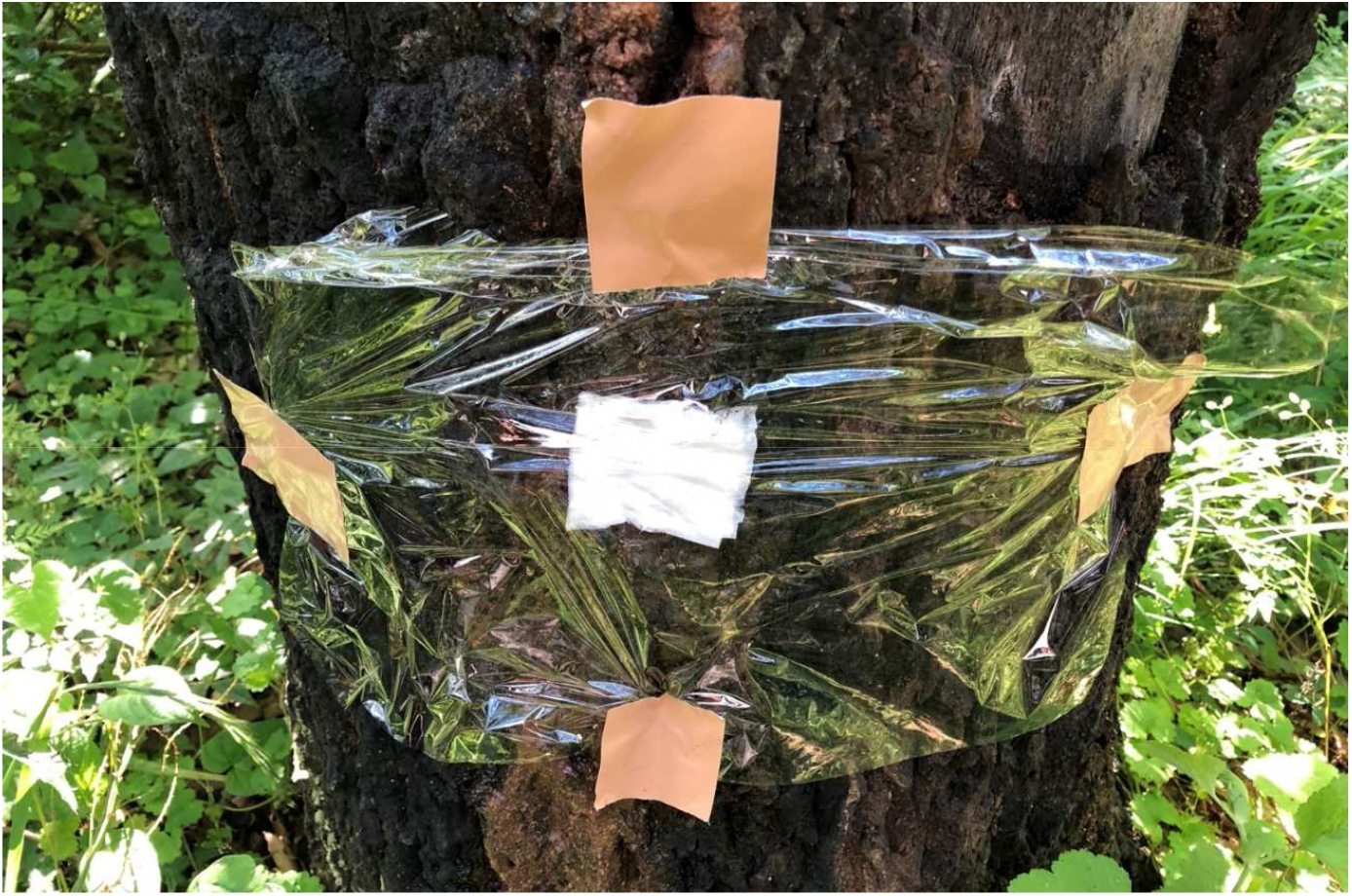
Experimental setup for measuring sap exuding rate on *Quercus acutissima*. The sap is absorbed by the paper towel at the center, and covered with cling wrap. Tape was applied to secure the cling wrap.

**Figure 3.**
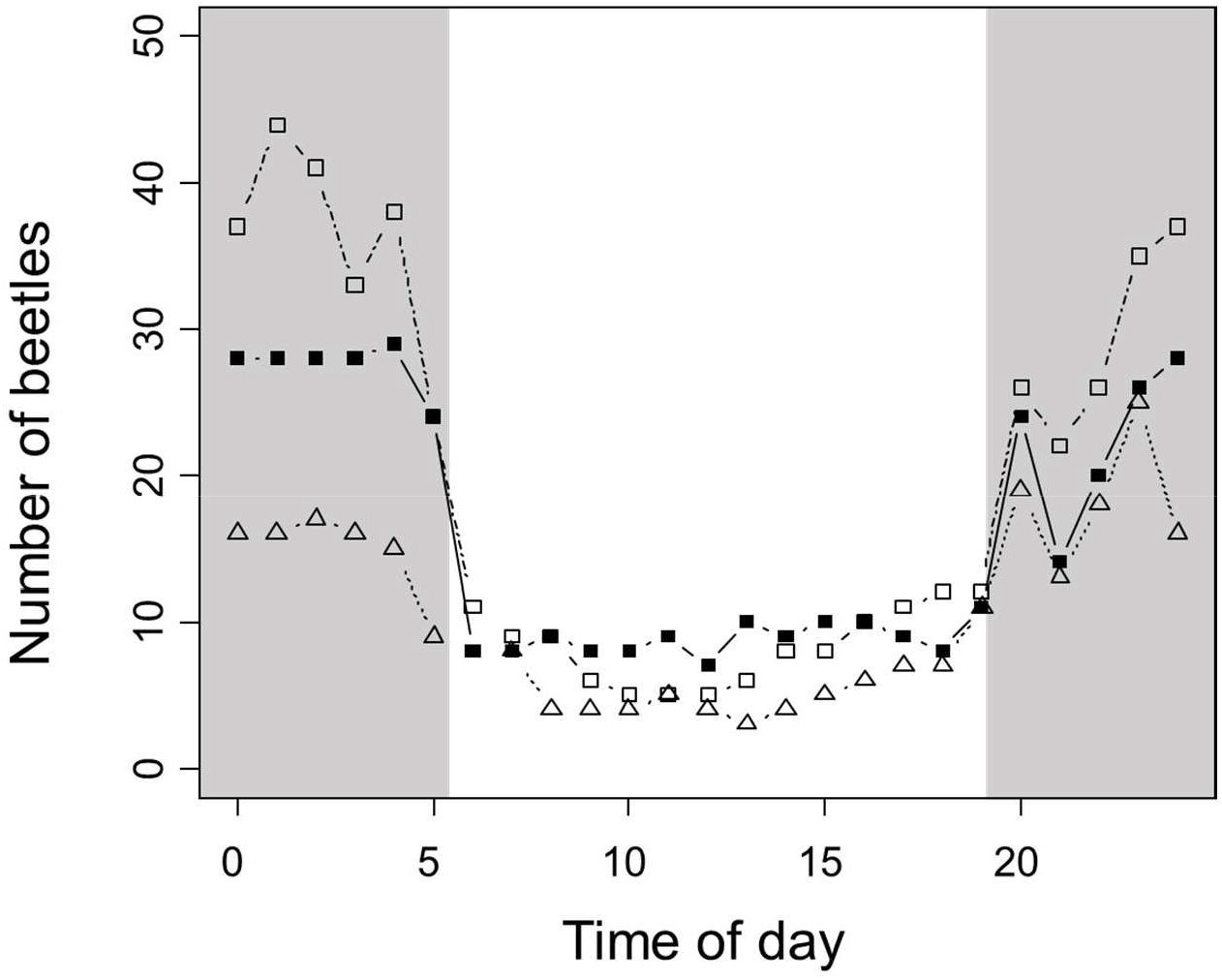
Daily activity pattern of *Trypoxylus dichotomus* on *Lagerstroemia subcostata*. The number of beetles is the total number of individuals in all surveyed trees. Different symbols represent the data of three days. The gray area indicates the night.

**Figure 4.**
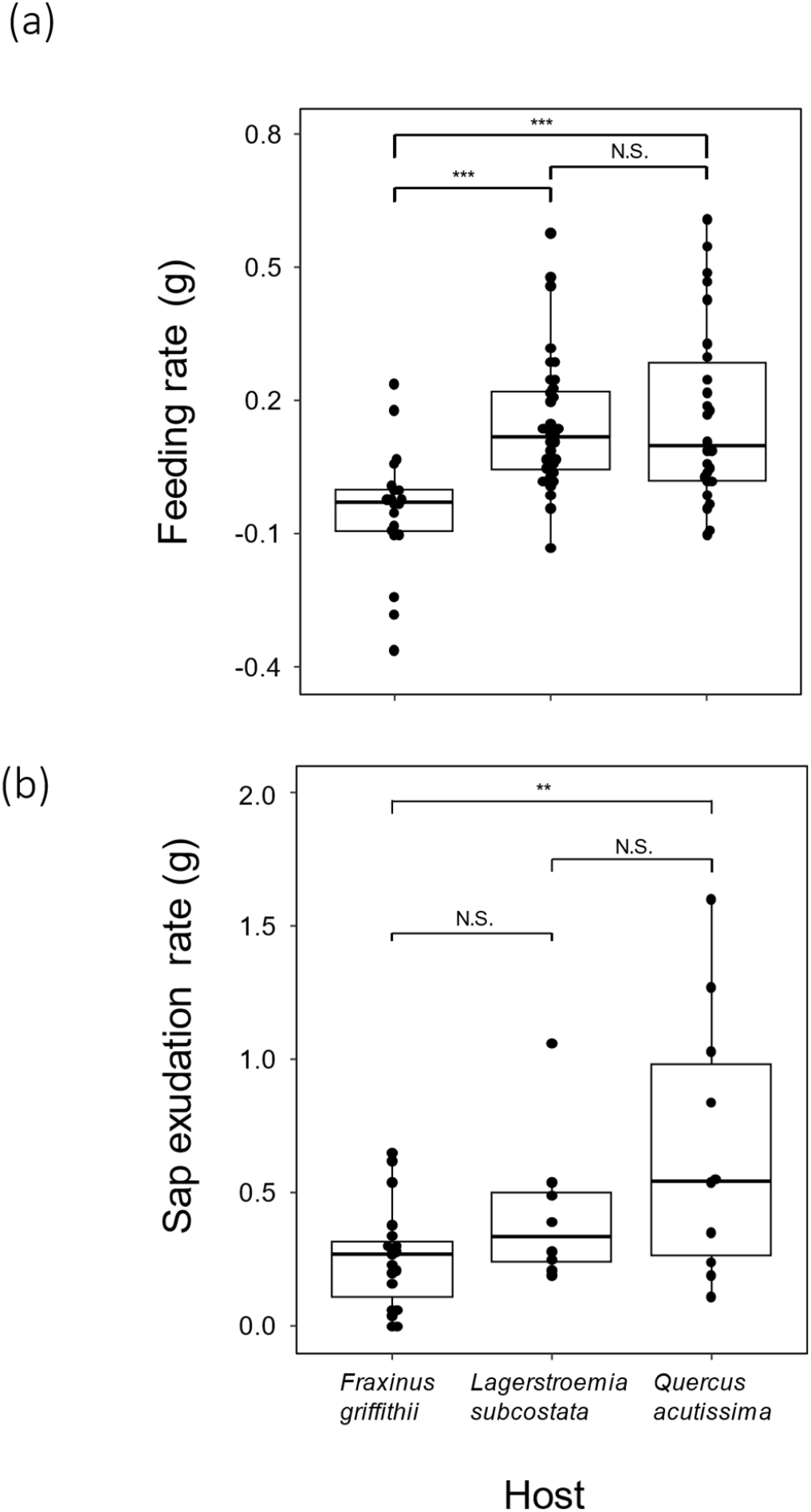
Feeding rates of *Trypoxylus dichotomus* (a) and sap exudation rates (b) on three host species. Feeding rates were assessed using hourly weight gain of the beetle. Sap exudation rates were assessed by measuring the weight gain during the 20-minute period that the paper towel was pressed against sap sites. Three asterisks indicate P < 0.001, two asterisks indicate P < 0.01, and ‘N.S.’ indicates no significant difference.

## Notes

### Competing Interest Statement

The authors have declared no competing interest.

### Summary of Updates

Added information in the Results section ; added information in the Discussion section ; added explanation of Figure 3 ; corrected unsupported references.

